# Establishment of microbial strains from the river, groundwater, and soil in the hyporheic zone is limited despite connectivity

**DOI:** 10.64898/2026.06.04.730025

**Authors:** Susan Mullen, Jacob West-Roberts, Lin-Xing Chen, Michelle E. Newcomer, Jordan Hoff, Eoin L. Brodie, Kenneth H. Williams, Shufei Lei, Jillian F. Banfield

## Abstract

Microbial diversity across ecosystems may be impacted by dispersal pathways and selection due to physiochemical conditions, and so it is hard to predict the extent of strain sharing across heterogeneous environmental compartments. Evaluation of patterns of strain-level overlap requires genome-level comparisons using extensive datasets collected over large spatial scales. Here, we sampled microbiomes beneath (hyporheic zone) and within the East River (Colorado, USA) and found little or no strain overlap at sites along the river corridor, despite connection by river flow, suggesting selection due to the local environment. Comparisons involving microbiomes from hillslopes and riparian zone soil yielded essentially no strain-level overlap with the microbiomes of river and hyporheic zones. We also sampled a nearby groundwater well and found near-perfect genotypic overlap with strains in one hyporheic zone location that exhibited active groundwater upwelling. Given an absence of direct connectivity between these two locations via discrete hydrologic flow paths, we conclude that groundwater strains are widely dispersed in the aquifer. As hyporheic zone microbiomes in zones with low river flow share no strains with groundwater microbiomes, we infer that strains introduced by groundwater mixing in the hyporheic zone are transitory. We conclude that despite evidence for mixing of river-and-hyporheic zone water, and river-and-groundwater on short time scales, establishment of transported strains in the hyporheic zone is minimal.

## Introduction

It is estimated that rivers and streams cover only half a percent of Earth’s non-glaciated terrestrial surface [1], yet they provide Earth’s population with over two-thirds of its drinking water. Rivers connect terrestrial and marine environments and are sites of extensive biogeochemical cycling. Water flows through the river channel and the porous space beneath the riverbed, known as the hyporheic zone, where river and groundwater mix. The interface between the river and hyporheic zone is dynamic, and groundwater mixing modulates geological, ecological, and biogeochemical processes [2]. Active weathering of the sediment releases electron donors and acceptors available to support microbial energy generation.

Prior studies of hyporheic zones have reported microbial processes, including heterotrophic aerobic respiration [3], anaerobic denitrification, dissimilatory nitrate reduction to ammonia [4], and nitrification [5,6] that play crucial roles in transforming a substantial portion of terrestrial organic matter [7–9]. Some studies on microbial communities of the hyporheic zone focused on biofilms on sediments [10], in pore water [11], and on hyporheic zone samples that contain both water and sediment [12], and most used 16S rRNA gene surveys [10–13]. These studies have brought to light the associations between microbial communities and grain size and geological features of the sediment [14], but offer an incomplete view of these complex ecosystems. More recent efforts have been made to classify the river water itself with metagenomic profiling [15], and a recent study focused on anthropogenic stressors across the hyporheic zone in North America, focusing on the sediment [16].

The East River originates north of Crested Butte, CO, in a mountainous watershed, and upon merging with the Taylor River, forms the headwaters of the Gunnison and Colorado Rivers **(Figure 1A)**. With an average elevation of 3266 m, the East River receives considerable snowpack over the winter months, with snowmelt in spring leading to substantial discharge through the river corridor [17]. In the summer, the discharge of the East River declines, and river water flows into the hyporheic zone. Previous metagenomic studies have been conducted on microbial communities in meander sediments flanking the East River [18] and hillslope soils and weathered rock [19]. The microbial communities of soil beneath the snow have been analyzed with metagenomics and metatranscriptomics [20]. East River hyporheic zone microbiomes have been studied at one site using 16S rRNA gene sequencing [21], but microbial metabolic capacities and lifestyles were largely unresolved. Hyporheic zone microbiomes are expected to change along the river corridor and with river discharge, likely impacting the processes that they mediate.

**Figure 1.**
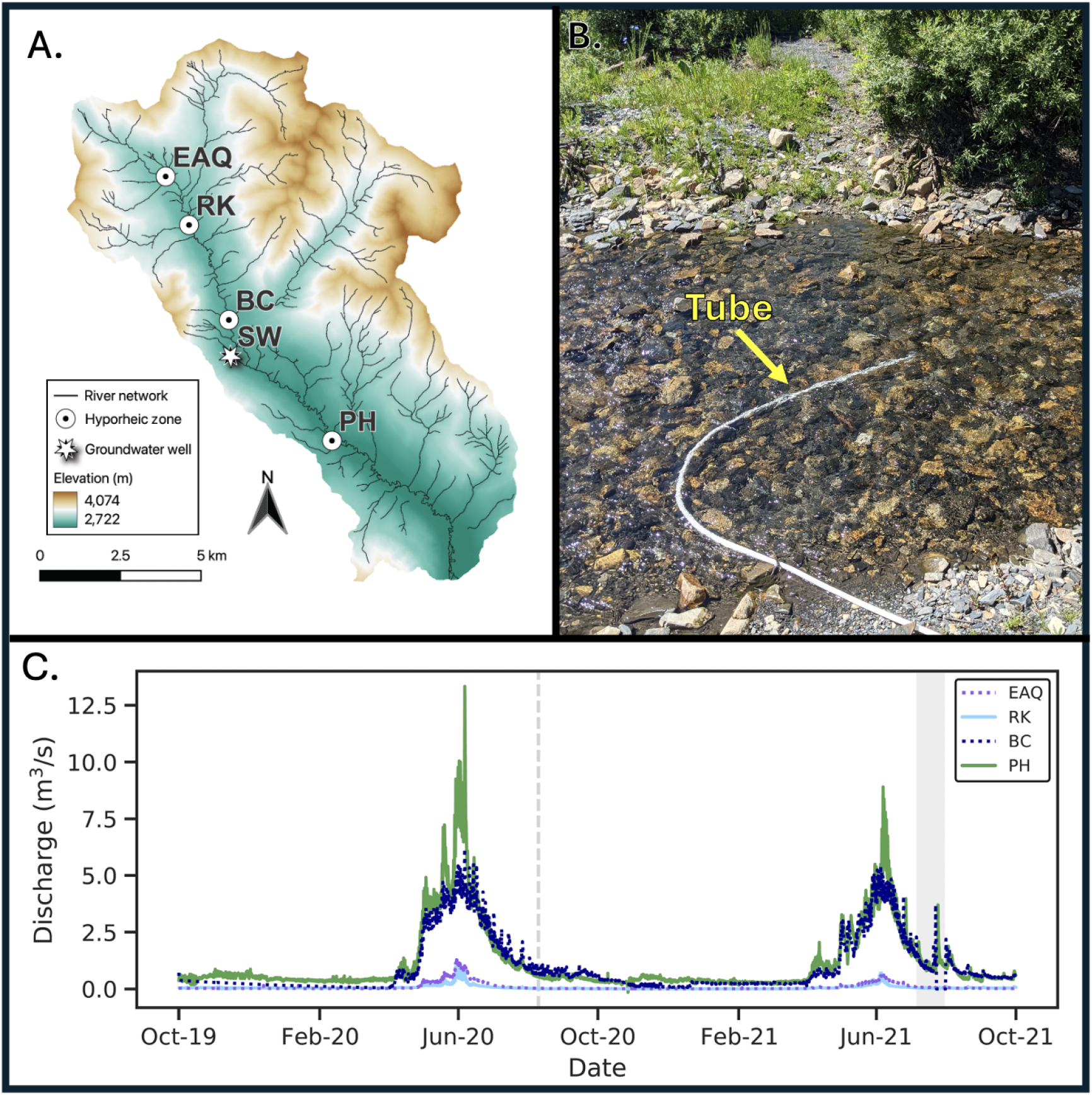
Sampling sites and temporal discharge: Sites are sampled in the East River, in Crested Butte, CO, USA. **A**. A map of the East River watershed made with Quantum Geographical Information System Version 3.40.5-Bratislava [61], displays the river. The streams shown in black are from the USGS National Hydrography dataset. The shape file was taken from [62]. Five sites were sampled: Shumway well (SW), East Below Copper (BC), Pumphouse (PH), East Above Quigley (EAQ), and Rock (RK). The hyporheic zone of BC, PH, and EAQ and the SW were sampled once in 2020. The hyporheic zone and river water of EAQ, PH, and RK were sampled multiple times in 2021. **B**. Field photo of the EAQ subsite B. The tube (20 mm external diameter) was buried in the center of the river to sample the hyporheic zone. **C**. Discharge flow of the river water at the site over a two-year period. The dashed line indicates when samples were collected in the summer of 2020, and the shaded area shows the timeframe when we sampled in the summer of 2021. The log scale of discharge is displayed in **Supplemental Figure 1**.

Here, we studied microbiomes of the hyporheic zone and associated river water at three sites along the East River at three time points as well as a groundwater well in the same watershed. We used genome-resolved metagenomics, which provides simultaneous insights into the composition of microbial communities as well as the metabolic potential of the community members. We hypothesized that, during periods of high river flow, hyporheic zone microbial communities would be more strongly influenced by river microbiomes than during low flow periods. Thus, we sampled over a three-week period in which river discharge decreased following snowmelt (falling limb). To introduce the possibility of differences in site-specific environmental conditions and to investigate hyporheic zone microbiomes that experience a range of river discharge rates, we sampled the river and hyporheic zones at sites up to 10 kilometers apart. Of particular interest were the questions of the extent of similarity between river and hyporheic zone microbiomes, the degree of overlap of microbes in microbiomes along the river corridor, and organismal overlap between the hyporheic zone and meander and hillslope soils.

## Results

### Study site and sample collectison

Although the East River is underlain by crystalline granitic rocks and hydrothermally altered sandstones of varying ages at higher elevations, the bedrock transitions to the late Cretaceous mudrocks of the Mancos shale along the river corridor, with this formation widely distributed throughout the central Rocky Mountains of the United States [22,23]. We selected study sites within the river corridor that are underlain by Mancos shale. The site at ∼ 2987 m on the main stem of the East River is referred to as the East River-Above-Quigley Creek (EAQ) site, which we sampled in 2020 and 2021. The Rock Creek (RK) site (∼2937 m elevation) is on a tributary of the East River, located ∼190 m from the junction with the main stem, and was sampled in 2021. Consequently, river water passing through the RK site is not directly inherited from the EAQ site. The Pumphouse (PH) site (∼2767 m elevation), is on the main stem downstream from both the EAQ and RK sites (**Figure 1B**). In 2020, we also sampled at the East River-Below-Copper Creek (BC), a site midway between EAQ and PH, which is on the main river corridor but has significant tributary input from Copper Creek, a stream within the Upper East River watershed.

Study sites were co-located with four concentration-discharge monitoring stations that were initiated in 2014 (PH) and 2016 (all other sites) and monitored at daily to weekly intervals over the period of sampling for this study. Notably, the downstream PH site experienced dramatically higher flow prior to and during this period than the EAQ and RK sites (**Figure 1C**). The RK site received less river water during the sampling period than the upstream site, EAQ. We sampled the hyporheic zone at the EAQ, BC, and PH sites at one time point in 2020. The BC site was replaced by the RK site in 2021 due to sampling difficulties, the onset of beaver activity, significant changes in stream flow behavior, and low levels of sequencing read assembly. Both the hyporheic zone and river water were sampled in 2021 at the EAQ, RK, and PH sites. At EAQ and RK samples were collected from three locations separated by ∼ 15 m at each site. These locations are named A, B, C. Note that we use the term site to represent EAQ, RK, PH, BC, and location to indicate the multiple closely spaced locations at RK and EAQ. The EAQ-C sampling location is close to the riverbank. At this site, we observed upwelling through the riverbed, despite downwelling being more common at this time of year. The EAQ and RK sites were sampled three times in 2021 over a three-week period, but the PH site was sampled only twice in 2021 due to a lack of site access. For hyporheic zone sampling, water was extracted from ∼30 centimeters below the sediment surface after the system recovered from disturbance associated with insertion of tubing (see Methods).

### Community composition analysis

We collected 23 hyporheic zone and 8 river water samples onto 0.1 µm filters and extracted whole microbial community DNA from which we generated 880 Gbp of raw metagenomic read data. Sequencing data were assembled into 170 Gbp of contiguous sequence (i.e., contigs). Of these, only 13% of assembled sequences were greater than 1 kb in length, likely indicating a wide diversity of organisms present at very low abundance levels. To profile the microbial community composition, we recovered the sequences of the ribosomal protein S3 (i.e., rpS3) and taxonomically classified each sequence with a database we made from rpS3s of GTDB genomes. Sequences were reconstructed from members of 63 bacterial and archaeal orders (**Figure 2**).

**Figure 2:**
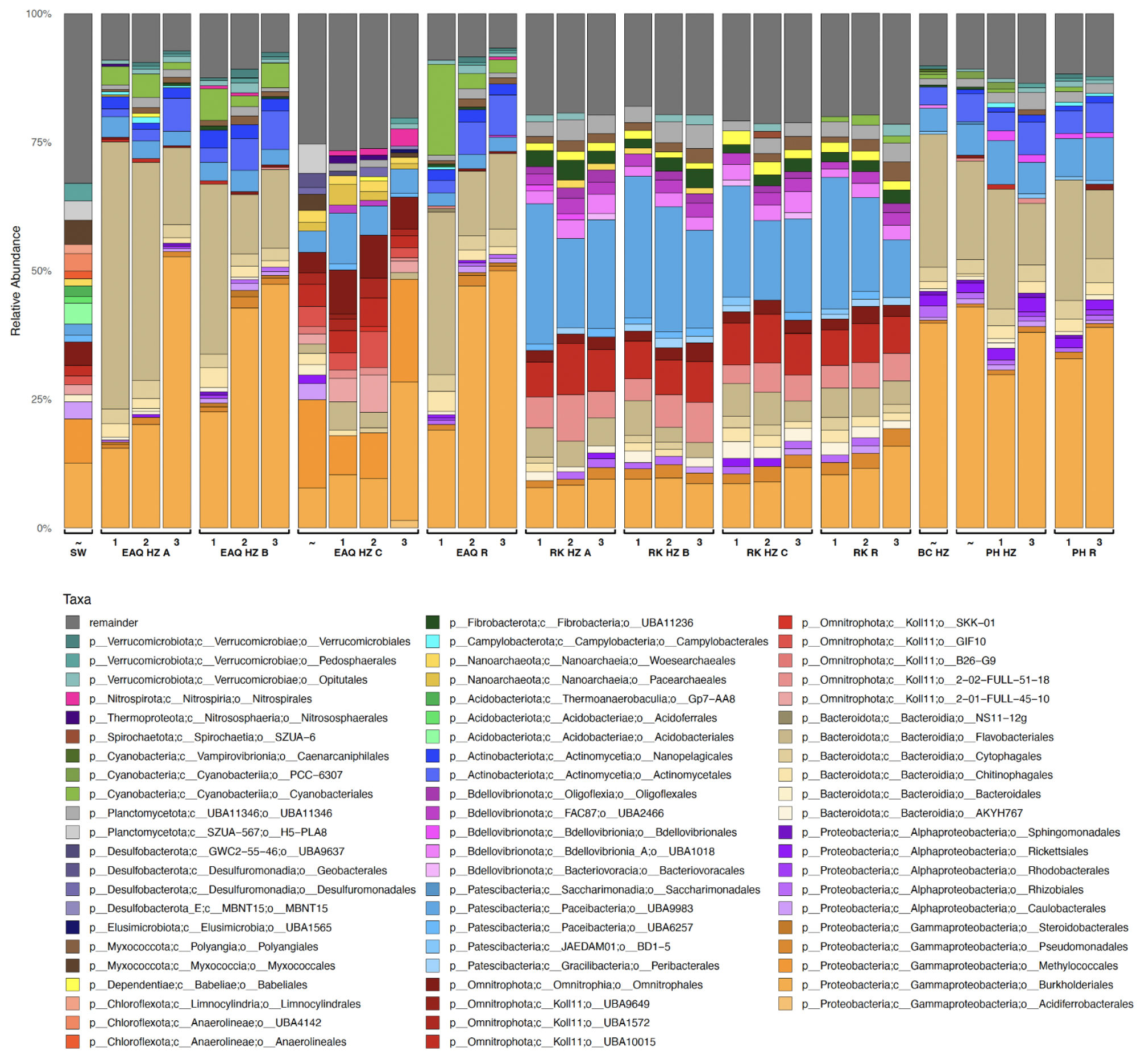
Community composition of hyporheic zone, river samples, and groundwater: The stacked bar graphs provide overviews of community composition at the order level. The ribosomal protein S3 sequences were used as the taxonomic marker for organism classification. The top 20 most abundant species for each sample are included; less abundant organisms were grouped into “other”. All organisms in the other category were present at < x% abundance level in the sample and can be found in **Supplemental Table 2**. R indicates river. The numbers indicate what week of sampling the sample was taken in July 2021, ∼ indicates that the sample was taken in August of 2020.

Across all sites, we detected Proteobacteria, Bacteroidetes, and Candidate Phyla Radiation (CPR) bacteria. Members of Methylococcales were notably more abundant than Burkholderiales in the EAQ-C river bank sample, whereas in all other samples, Burkholderiales was the most abundant group. In some sites, we detected bacteria assigned to Omnitrophota, Bdellovibrionota, and Myxococcota.

Regardless of whether the sample was collected from the river or from the hyporheic zone, the microbial communities cluster into three community types **(Figure 3)** delineated at the order level **(Figure 2)** and in a dendrogram calculated using the relative abundance data **(Figure 3)**. Type 1 includes all three 2021 hyporheic EAQ-C samples and the 2020 EAQ hyporheic zone sample from the same location. Unlike the other types, Type 1 microbiomes have a high relative abundance of three uncultivated Omnitrophota: GIF10, UBA1572, and JAHIP101.

**Figure 3:**
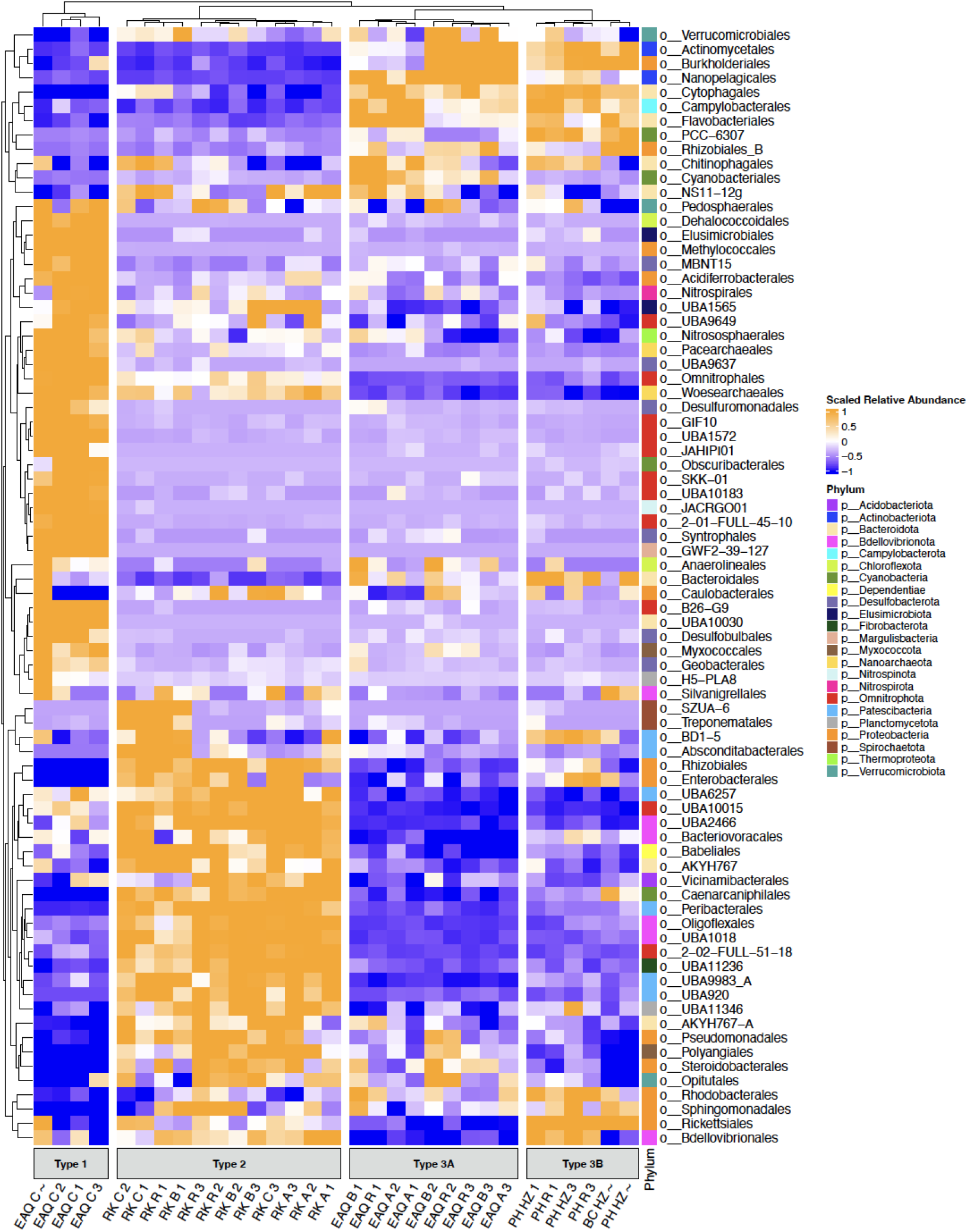
Type 1, 2, and 3 community compositions in the hyporheic zone and river based on relative abundance patterns: Heat map showing relationships between the community composition of the relative abundance: orange represents high scaled relative abundance and blue represents low. Data were hierarchically clustered to show relationships across samples. Two main cohorts emerge. RK forms cluster 2. Pump House, EAQ, and BC form their own cohort. Copper is most similar to Pump House. EAQ-C has its own unique community composition and microbial signature. The numbers indicate what week of sampling the sample was taken in July 2021, ∼ indicates that the sample was taken in August of 2020.

Type 2 microbiomes include all of the RK site (river and hyporheic zone) samples. They have high levels of another uncultivated Omnitrophota, 2-02-FULL-51-18, which are rare or not detectable in the other microbiome types.

Type 3 microbiomes have low abundances of Omnitrophota and are divided into two groups based on overall microbiome composition. Type 3A are the EAQ sites exclusive of EAQ-C, and Type 3B are the PH and 2020 BC sites.

The EAQ-C (Type 1) samples provide insight into riverbank hyporheic zone microbial communities, as they were the only hyporheic samples not collected from the center of the river. In contrast, the other EAQ site hyporheic zone microbiomes, and the PH hyporheic zone microbiomes resemble those of the respective river samples, likely indicating that the hyporheic zones at the center of the river at this time of year are strongly impacted by downwelling river water. The species or higher taxonomic level dissimilarity between the RK river microbiome and other river microbiomes and similarity between RK river and RK hyporheic zone is most likely due to the differences between the tributary and river, as RK is the only site located on a tributary.

### Organism distribution and dispersal within the hyporheic zone and river sites

To understand microbial distribution patterns, we identified groups of similar organisms by constructing phylogenetic trees for organisms from six phyla using concatenated ribosomal protein sequences (**Supplemental Figure 2**). Mostly, we found the same species at different locations (e.g., A, B, or C) within a specific site, although some similar species were detected in different sites. To more stringently identify organisms shared between sites, we clustered genomes based on 99.5% average nucleotide identity (ANI) and used these results to look for evidence of strain sharing along the river. In this system, dispersal should be facile given site connectivity by river flow. Thus, we asked whether organisms dispersed across sites are recruited into microbiomes or selected against. We found evidence for the sharing of strains abundant enough for genomic reconstruction in very few samples from different sites or site types. Three strains are shared by river and/or hyporheic zone samples from different sites, and one strain is shared by hyporheic zone samples from two different sites (**Supplemental Table 1**).

### Organism overlap across the hyporheic zone, hillslope transect, and meanders

Watershed and hillslope hydrologic connectivity may provide the opportunity for organism overlap between different regions of the watershed, although environmental conditions may select against dispersed strains. To investigate this, we compared the organisms detected in metagenomes from (a) three hyporheic zones (EAQ, RK and PH) and river to those of previously studied floodplain meander soils (meander, G sampled close to EAQ, meander L, which is adjacent to the PH site and meander Z; Matheus Carnevali et al. 2021) [18] and (b) five soil sites along a hillslope adjacent to the PH site (Lavy et al. 2019) [19]. In total, the dataset included 145 metagenomes. Eleven datasets, comprised of three hyporheic zone datasets, three meander datasets (G, L, and Z), and five hillslope datasets (PLM0, PLM1, PLM2, PLM3, PLM4), with all replicates and time points combined, were compared in a pairwise manner using rpS3 gene sequences as organismal markers.

We took datasets that included the river and hyporheic zone sequences from each site and compared them with those of the meander soil sites. First, we identified taxa as shared if their rpS3 genes had nucleotide identity of >90%. We noted both the number of sequence pairs that met this identity threshold and the taxonomy of each organism pair (**Figure 4**). The RK site shares the fewest organisms with all of the meander sites, possibly reflecting the location of the RK site on a tributary. The meander sites share more organisms with EAQ than the other sites. The PH site shares many organisms with the upstream meander G and co-located meander L, yet only two organisms are shared between the PH and the downstream meander Z site.

**Figure 4:**
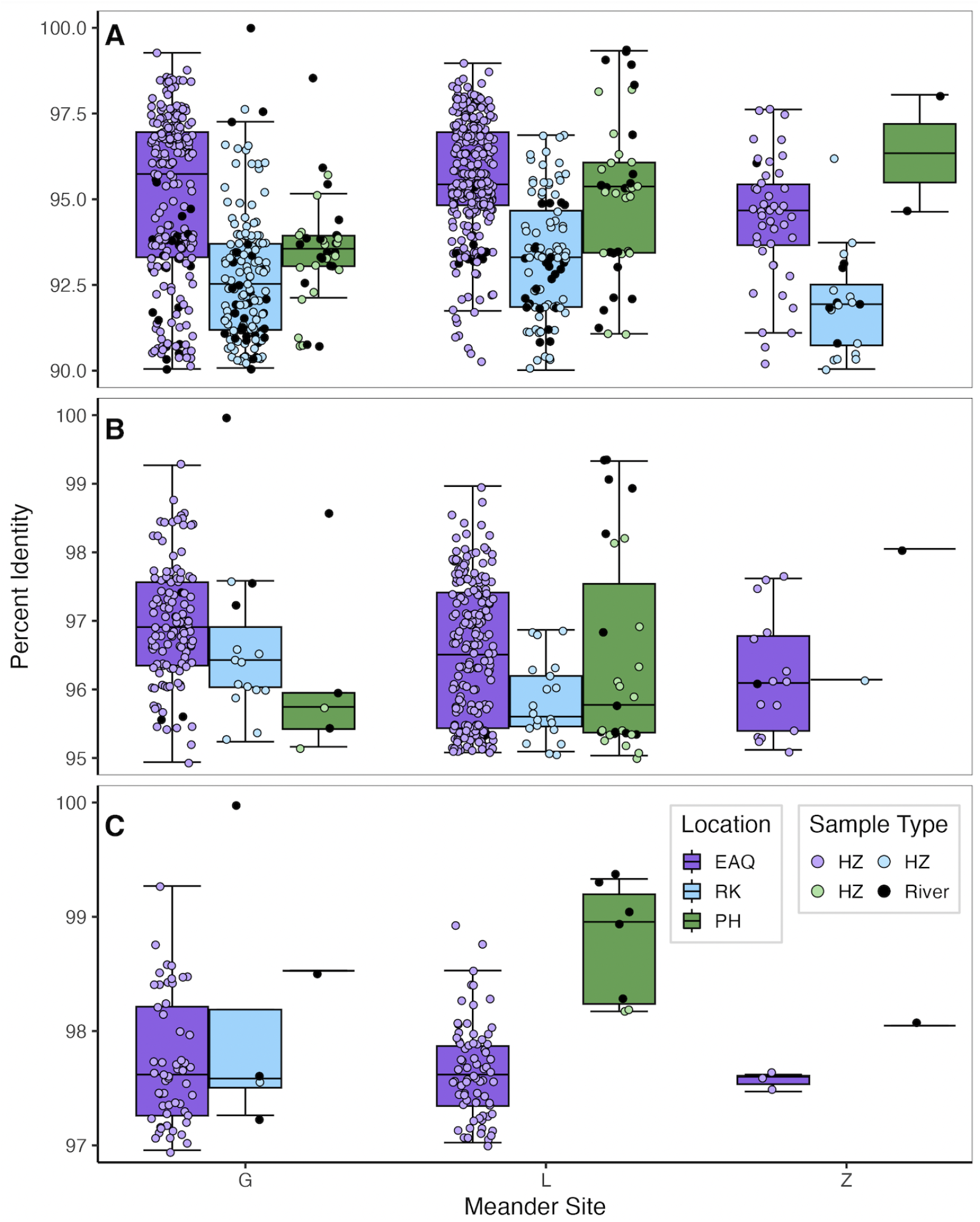
rpS3 comparison of three meander-bound floodplains in the East River meanders to the hyporheic zone and river sites. Box and whisker plots showing the percent identity with which the rpS3 genes of the hyporheic zone and river dataset align with the rpS3 genes of the hillslope dataset. **A**. Organisms that share 90-100% nucleotide identity. **B**. Organisms that share 95 - 100% nucleotide identity. **C**. Organisms that share 97-100% nucleotide identity. The colors purple, light blue, and green, represent the location of the river and hyporheic zone data, EAQ, RK, and PH, respectively. Black filled points represent the alignment of a meander rpS3 gene with an rpS3 gene from a river sample. Whereas blue, green, and purple filled points represent overlap with an rpS3 gene from a hyporheic zone sample. The boxes are grouped by meander location. Meander L and PH are co-located.

Next, we investigated the extent of overlap of more closely related organisms using thresholds of > 95 % and > 97% rpS3 nucleotide identity, as described in Olm et al. 2020 [24]. At the > 97% rpS3 approximately species level, the upstream EAQ site shares species with all three downstream meander sites. The PH site shares some species with its associated meander L, but only one with meander Z and one with meander G. There is almost no species overlap between the downstream meander Z site and the PH and RK sites. As only one species shares >99.5% rpS3 nucleotide identity (meander G and rock site river) we did not pursue genome-wide comparisons to look for strain sharing between the meanders and the hyporheic zone plus river datasets.

We conducted a similar comparison involving samples from the hillslope sites. At the >95% threshold, we found very little overlap between the RK site and any of the hillslope sites, and at the >97% threshold, there is no overlap **(Figure 5)**. At the 97% threshold, the EAQ hyporheic zone site only has overlap with the three hillslope sites that are closest to the river’s edge (PLM4, PLM3, and PLM2). At this threshold, the PH site also shares organisms with river edge hillslope sites and shares one organism with a site that is higher upslope **(Figure 5)**. No pair of organisms shared >99.8 % rpS3 nucleotide identity, so we did not pursue genome comparisons to investigate sharing of strains between the hyporheic zones and hillslope.

**Figure 5:**
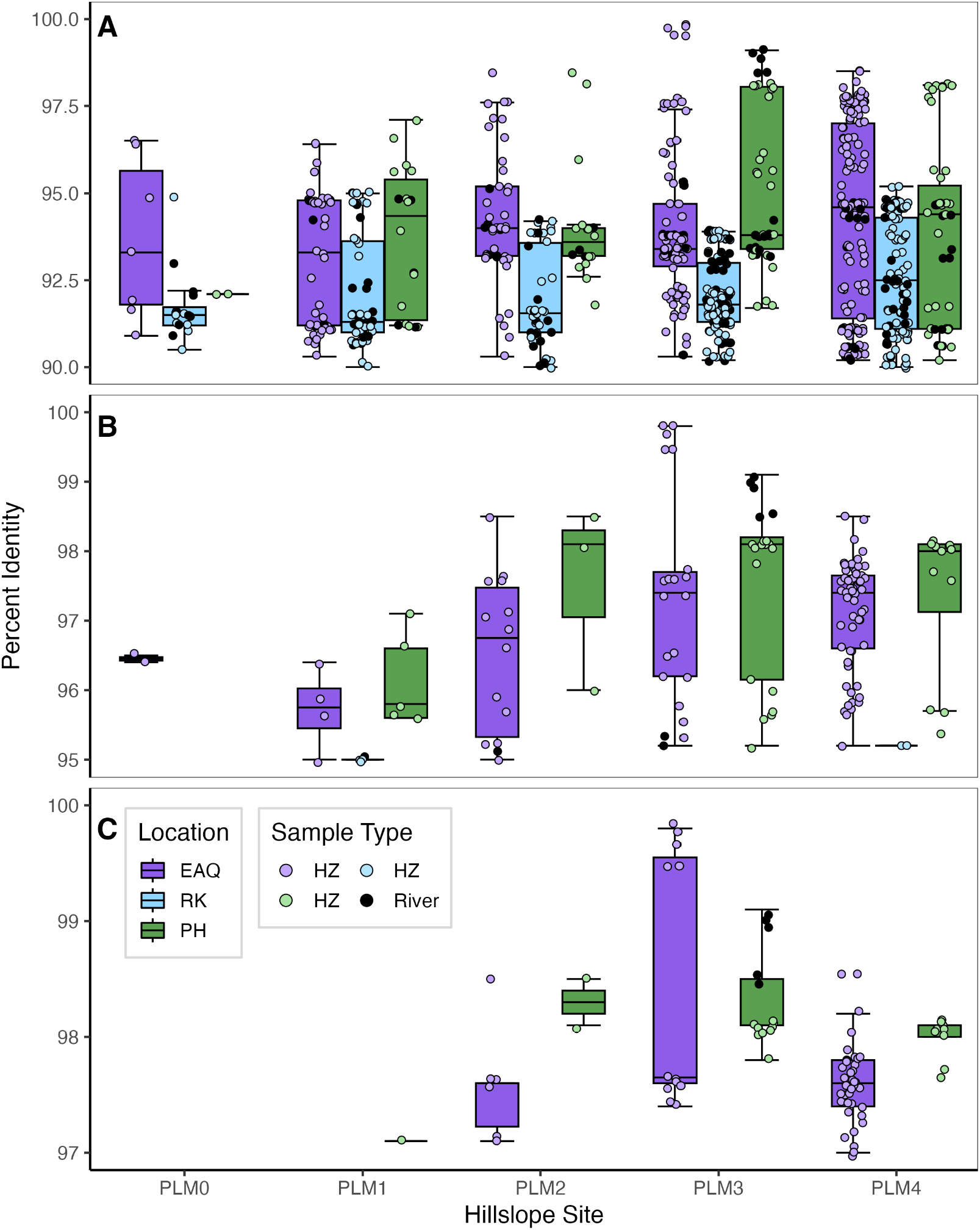
rpS3 comparison of hillslope transect in the East River to the hyporheic zone and river sites. Box and whisker plots showing the percent identity with which the rpS3 genes of the hyporheic zone and river dataset align with the rpS3 genes of the hillslope dataset. The colors purple, light blue, and green, represent the location of the river and hyporheic zone data, EAQ, RK, and PH, respectively. The boxes are organized by the hillslope transect they align with. Black filled points represent the alignment of a hillslope rpS3 gene with an rpS3 gene from a river sample. Whereas blue, green, and purple filled points represent overlap with an rpS3 gene from a hyporheic zone sample. The hillslope transects are physically closest to the PH site.

### Strain sharing between groundwater and the upwelling hyporheic zone site

We tested whether there is evidence for groundwater microbes in the hyporheic zone by comparing the hyporheic zone microbiomes with those of the aquifer, as represented by groundwater collected from the ∼70 m-deep Shumway well in nearby Mancos shale bedrock (**Figure 1A**). After clustering genomes from all hyporheic zone and river samples and the Shumway well at 95% ANI, only genomes from the one 2020 and three 2021 EAQ-C hyporheic zone samples were shared with those from Shumway well. This groundwater well is connected to the aquifer of the East River watershed, but it is not directly upgradient from any of the hyporheic zone sites. We performed a rigorous strain comparison between the EAQ-C and Shumway microbiomes using inStrain and considered draft genomes that shared > 99.999% population average nucleotide identity (popANI) to be from the same strain. We identified 14 Shumway and EAQ-C genome pairs that met this threshold (with ≥ 99% breadth of coverage of the respective bins). The 14 genomes represent bacteria from, Bacteroidetes (3), Ignavibacteria (1), Myxococotta (1), Omnitrophota (1), Planctomycetota (2), Proteobacterota (5), and Verrucomicrobiota (1). Thus, we infer substantial input of the regional groundwater microbiome, represented by the Shumway well, via upwelling into the EAQ-C hyporheic zone, rather than upwelling of locally sourced groundwater with a substantially different microbiome.

### Giant proteins and other genes potentially involved in inter-organism competition

Given the high abundance of Omnitrophota in the RK and EAQ-C sites and Shumway well, and the connection between Omnitrophota and giant proteins, we searched for giant proteins of > 30,000 a.a. We found two giant proteins in the EAQ-C sample, both of which are on contigs that were derived from Omnitrophota genomes. Manual curation extended the first gene to a stop codon, resulting in a 217,174 bp open reading frame coding for a complete 71,949 a.a. protein (**Figure 6A**). The protein has a low density of detectable PFAM domains (12.2% when using standard GA cutoffs), consistent with previous analyses of these proteins [25]. Observed domains include glycosyl transferases/hydrolases, glycolysis machinery, methyltransferases, and GGDEF c-di-GMP signaling domains (**Figure 6B**). Manual curation extended the second gene to 131,644 bp, representing a protein of >43,881 a.a. (curation could not extend to the stop codon). Unlike the first case, we were able to extend the second contig far enough to document upstream genes, which encode primarily for components of the type II secretion system proteins (**Figure 6C**).

**Figure 6:**
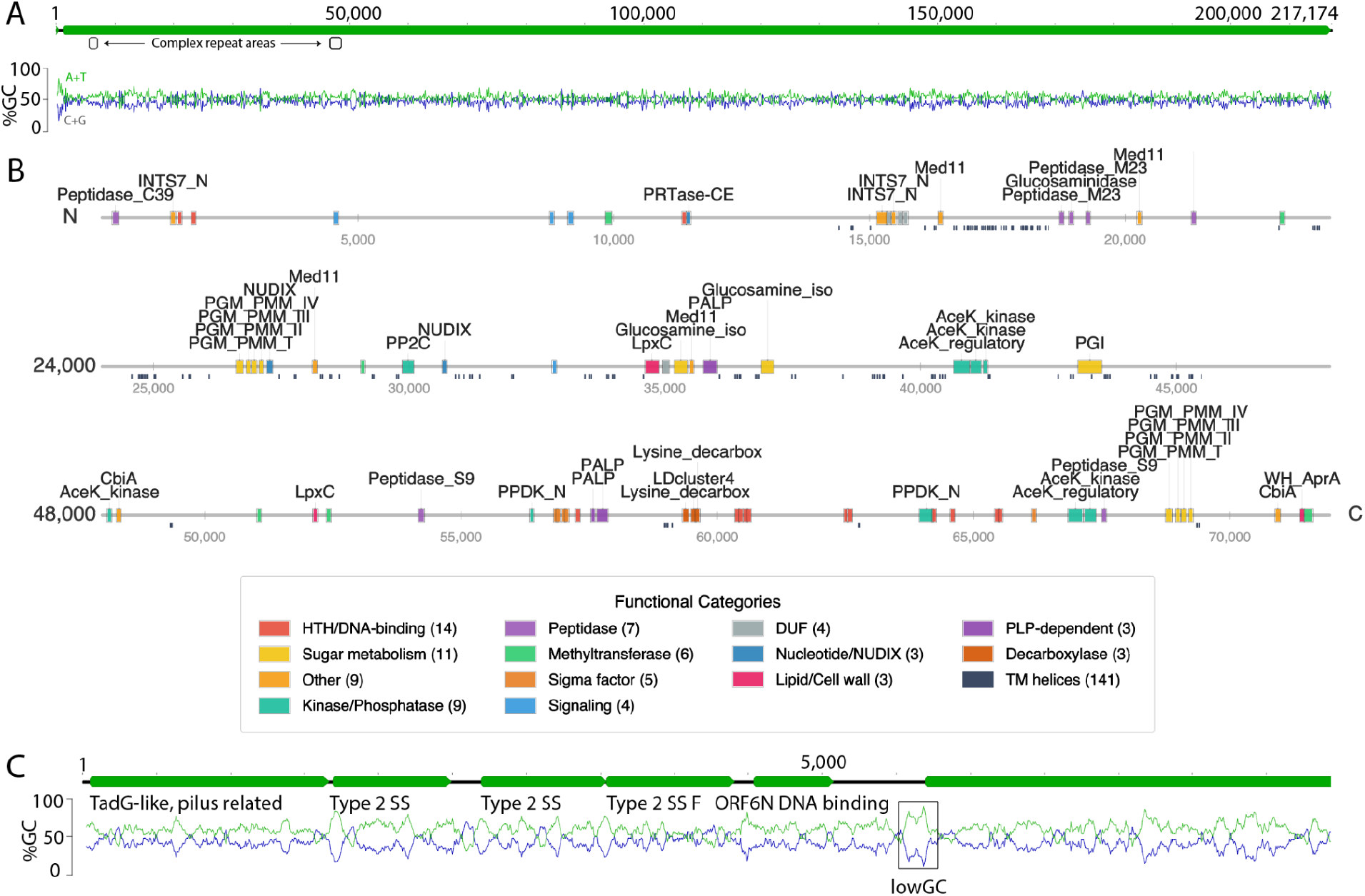
Giant proteins of Omnitrophota in the EAQ-C sample. **A**. Fragment of an Omnitrophota genome encoding a large partial open reading frame was curated to extend the sequence, resolve complex repeat regions (boxes) and remove scaffolding gaps. Recovery of the complete 217,174 bp gene is evidenced by preceding open reading frames and a stop codon. **B**. Diagram illustrating features of the giant protein in A. Notable are domains involved in protein and carbohydrate compound transformations, such as LpxC (UDP-3-O-acyl-N-acetylglucosamine deacetylase) involved in lipid A modification, PGK (phosphoglycerate kinase) /TIM (triose-phosphate isomerase) domains found in glycolytic enzymes, glucosamine_iso (glucosamine-6-phosphate isomerase) involved in aminosugar metabolism, PGM (phosphoglucomutase /phosphomannomutase) domains for glycan biosynthesis and degradation, glucosaminidase for glycoprotein degradation, INST7 domains resembling the Eukaryotic integrator complex, which cleaves nascent mRNA, and PRTase (phosphoribosyltransferase) that may function in nucleotide salvage. **C**. Genes that precede the start of the second large (> 131644 bp) Omnitrophota gene (stop codon not reached during curation) encode mostly secretion system-related proteins. The giant gene is preceded by a much lower (∼16%) GC-content region that lacks a detectable Shine–Dalgarno ribosome-binding site and recognizable σ^70^ promoter elements, suggesting the use of atypical or internal promoters and potentially leaderless translation initiation.

Given the prevalence of Omnitrophota, Bdellovibrionota, and Myxococcota, organisms previously implicated in predation [e.g., 26–28], we tested for complexes (in addition to the giant proteins) that may be associated with antagonism, such as RHS toxins, type VI secretion system (T6SS) and T6SS effectors, and lysis machinery (**Supplemental Table 3**). We acknowledge that these systems may also be involved in microbial processes such as defensive weapons, nutrient acquisition, pathogenesis. Most prevalent were T6SS, RHS toxins and lytic enzymes.

## Discussion

We integrated metagenomics and spatial analyses primarily to investigate microbial community structure, dispersal, and adaptation in the hyporheic zone and water of the East River. To evaluate the extent of engraftment of dispersed species, we compared the species and strain composition of the hyporheic zone and river with the hydraulically connected hillslope, three floodplains, and a nearby groundwater well. In the summertime, the microbiome of hyporheic zone samples collected at the center of the river, reflect the microbiomes of the river water samples at their respective sites. The river water microbiome at the RK site has a different overall microbiome composition than the river water at PH, EAQ, and BC sites, likely because the RK site is on a tributary that is known to drain from an organic matter rich peatland. In the summer, the RK river water has a distinct dissolved organic carbon signature that is higher than those of the other sites **(Supplemental Figure 3)** [29]. The EAQ-C site, which was collected at the bank, is more similar to the RK site, probably because bank sediment is typically richer in organic carbon than sediments from the center of the river [30,31].

We observed only a few instances of species overlap involving the hyporheic zone and the river at different sites along the river corridor. This may be due to site-specific properties at the EAQ, RK and PH sites that impose niche constraints. Our analyses indicate that hillslope and floodplain microbiomes share more species overlap with each other than with the river and hyporheic zone, likely due to similar environmental conditions. Dispersal is enabled by translocation of eroded hillslope materials (e.g., soils, sediments, bedrock fragments) to the floodplain. The microbes of the tributary RK hyporheic zone and river have no discernible impact on downstream hyporheic zone and river site microbiomes. We suggest that, despite the potential for river transport, the microbes of the RK site river and hyporheic zone are not adapted to the conditions at the other sites. This is in agreement with the Graham et al. 2017 study, which showed that while dispersal has some impact on microbiome community composition in a river ecosystem, selection is the main driver of microbial establishment [32].

In cases where marker gene sequence similarity was very high, we used genomes to accurately test for organismal overlap. The thresholds of >99.999% ANI over 99% of the genome enable strain tracking. Overlap involving 14 strains shared by the Shumway groundwater and the EAQ-C sites establishes microbial input from upwelling groundwater. The groundwater likely also mixes with river water in the other hyporheic zone locations, as the hyporheic zone, by definition, receives a mixture of river water and groundwater. As all other hyporheic zone sites did not show strong upwelling and do not share strains with the groundwater, we infer that the strains introduced by the influx of groundwater to the hyporheic zone are transient. The effect at EAQ-C may have been reinforced by the low river flow rate at the time of sampling. The high river flow rate at the PH site at the time of sampling likely resulted in the river microbiome overwhelming any hyporheic zone adapted strains and strains from mixing with groundwater. We conclude that generally the aquifer microbes do not persist in an abundance sufficient for detection in most hyporheic zone regions, at least over the late summer.

The giant proteins observed in Omnitrophota share similar features to those previously described in Omnitrophota genomes [25,28,33]. Related Omnitrophota proteins were also detected in the PH and Shumway well microbiomes, but the sequences were too fragmented for further analysis. The giant proteins have functional domains for carbohydrate metabolism, lipid synthesis, and nucleotide salvage. Domains for carbohydrate metabolism and lipid synthesis suggest roles in cell surface modification. Some domains, including peptide digestion and carbohydrate metabolism, may be involved in the breakdown of cell walls of other organisms, consistent with experimental studies that implicate Omnitorophota in predation [28,33]. The Type VI secretion system enzymes, toxins, effectors and lytic enzymes may also play roles in lethal competition.

Based on a study of microbiomes sampled throughout the East River watershed, we conclude that organic carbon, influx of regional groundwater, and hyporheic zone conditions shape the community in and beneath the river corridor. The magnitude of each factor varies substantially in its impact on river and hyporheic zone microbiomes, and results in little species overlap, despite broadly similar habitats at different sites along the river. In a study of the Columbia River and associated hyporheic environments within and at some distance from the river, Graham et al. (2017) found that spatial variation in hyporheic zones imposes strong selection effects that limit the persistence of microbes introduced from other sites. We found that microbial species and strains sourced from the hillslopes and floodplain soils are not readily detectable in the river and hyporheic zone. Notably, genomic-level analyses of strain overlap confirmed that organisms introduced by upwelling groundwater (and apparently representative of regional groundwater) do not establish in the hyporheic zone at detectable abundance levels. Although there is connection along the watershed between the river, groundwater, and soil of the hyporheic zones, strains are not widely shared or established beyond their local environments.

## Materials & Methods

### Site Description & Sample Collection

Sites selected were historical concentration discharge sampling sites, for which there were existing water chemistry and discharge data collected over multiple years [34,35]. A discharge data package is available for Pumphouse (PH), Rock (RK), East Above Quigley (EAQ), and East Below Copper (BC) [36]. We chose three main sites to represent upstream, downstream, and midstream of the East River catchment. The most upstream site and midstream site were not connected to one another, but both the upstream and midstream sites are connected to the most downstream site. The downstream site was less accessible than the other two sites and we relied on transportation from staff scientists in an all terrain vehicle.

We confined our study to the period of decreasing flow (late summer) due to safety concerns associated with accessing the hyporheic zone during maximum flow following snowmelt (spring to early summer). We did not sample during the lowest flow periods (winter) as the site would be covered in snow (causing our peristaltic pumps to freeze). Due to the limitations of accessibility, we did not sample the Pumphouse site as often as the other two sites, and we did not install replicate hyporheic zone samples. All but one sampling location had a rocky upper layer.

Beneath that layer was gravel and sediment. At the sampling locations, we dug 30 centimeter deep holes and buried one tube per hole with the opening of the tube facing downward. The 30 centimeters was measured from the start of the sediment layer. All but one sample was located equidistant from the riverbank.

We put a stainless steel mesh casing on the entrance of the tube to prevent larger rocks from getting sucked into the tube. There was one location, this site East Above Quigley-C, that was measured closer to the riverbanks and not equidistant from the riverbanks; it had a noticeable plume of upward movement of water. There were particles of fine sediment that were moving up the column of water that we did not see at any other site. EAQ-C did not have a noticeable rocky upper layer; it was primarily composed of gravel and sediment. EAQ site may offer variability in hydrological flow paths due to a beaver dam upstream and visible turbulence. We refilled the holes first with sediment and gravel that was dug up, and then we covered them with any rocks that were dug up. We waited at least 48 hours before beginning the sampling to let the disturbance of the digging settle and for the system to be as close to its natural state.

To filter river water from the horizontal and vertical center of the river, we weighed down a tube so that it was approximately halfway between the bed of the river and the surface of the river. We used a model 77602-10 masterflex easy-load peristaltic pump and masterflex 6424-82 tubing and filtered 100 gallons of water or until the filter clogged and no more water could be filtered. The filters used were Grover Technologies ZTECG 0.1 um - 5P3S. Tubes were sterilized in advance with 10% bleach and flushed with Milli-Q water. We set the speed of the peristaltic pump, such that the flow rate would start at 1 gallon per minute as measured by DigiFlow6700M. The flow rate did slow down as the sampling went on, this was likely due to sediment clogging the filter. We removed the filters with sterile gloves and stored them in WhirlPak bags. The samples were then put on dry ice and transported back to the University of California, Berkeley, and stored in a -80C freezer until they were extracted.

### DNA extraction and sequencing

Filters were thawed at room temperature for thirty minutes and cut with sterile pipe cutters to remove the plastic casing. One third of the filter was cut off with a scalpel for extraction, and the rest was saved in the -80C freezer. Qiagen DNA Power Soil Kits were used to extract the DNA from the filters, which contained cells from the water and cells attached to sediment-associated particles. The protocol was modified to accommodate the filter. The filter was vortexed with PowerBead solution for 10 minutes at high speed. Then the lysis buffer was added to the solution, and this mixture was incubated for 10 minutes in a 65C water bath with occasional mixing. Then the Qiagen protocol was followed and the bead was stored elsewhere [37] and stored in molecular grade water in the -80C freezer. Genomic DNA yields were between 10 ng/uL and 70 ng/uL, except for five samples, which were between 3 ng/uL and 5 ng/uL. Samples were diluted to 10 ng/uL in 50 uL. Those that had lower starting concentrations were not diluted. DNA was quantified using a Qubit double-stranded DNA high-sensitivity assay. Metagenomic libraries for all samples but Shumway were prepared and sequenced at the University of Maryland using an Illumina NovaSeq6000 SP platform to generate 250-bp paired-end reads, generating 23-25 Gbp per sample. Samples were multiplexed for sequencing. The Shumway sample was sequenced at UC Berkeley QB3 using an Illumina NovaSeq6000 SP platform to generate 250-bp end reads, generating 20 Gbp.

### Metagenomic analysis, assembly, and binning

Reads from individual samples were assembled using Metaspades Version 3.15.5 [38]. Only assembled scaffolds with 1000 bp were included in downstream analysis. Trimmed sequencing reads from each sample were mapped. Bins were obtained using automated binners Maxbin2 [39], MetaBAT2 [40], Concoct[41], and VAMB[42]. The best set of bins was selected using DAStool [43] as described by Diamond et al. [44]. This resulted in 1362 draft genomes. Open reading frames were predicted with MetaProdigal [45]. Predicted ORFs were annotated using KEGG [46], Uniprot [47], and Uniref100. These draft genomes were dereplicated based on 95% Average Nucleotide Identity (ANI) using DRep [48] with the –ignoreGenomeQuality flag. This resulted in 251 representative genomes. These 251 genomes were manually curated on our in-house analysis database ggKbase (ggkbase.berkeley.edu) based on visual inspection of taxonomic profile, GC content, coverage, and a set of 51 bacterial single-copy genes (BSCG), and 38 archaeal single-copy genes (ASCG) [49]. Genome coverage was calculated by mapping sample reads to the genomes with BBMap (v.39.10) and summarized using CoverM [50].

### Community composition and lifestyle analysis

We used ribosomal protein S3 (rpS3) as a marker gene for taxonomy and used KEGG orthology ID K02982 as the HMM for our query. Aksha, an HMM wrapper script [51], was used to identify rpS3 genes. rpS3 genes were clustered using MMseqs at 80%, and we selected the longest sequence to be the centroid. The nonredundant centroids were then mapped to the reads using BBMap, minid=0.95, idfilter=0.97. CoverM was used to assess the mean relative abundance of each sequence. References were chosen using BlastP and GTDB-TK [52]. Taxonomy up to the order level was assigned. We displayed the top 20 relative abundances for each sample in **Figure 2**. We used the abundance patterns of rpS3 and hierarchically clustered them using the Jaccard method **(Figure 2)**. The Jaccard method was used to cluster the samples based on abundance patterns, and the heatmap was produced with Complex Heat Maps[53] (**Figure 3**).

For further analysis, phylogenetic trees were constructed with a set of 16 ribosomal proteins: L2, L3, L4, L5, L6, L14, L15, L16, L18, L22, L24, S3, S8, S10, S17, and S19 after Hug et al [54]. Homologous ribosomal proteins were aligned using MAFFT (version v7.505) [55], and trimmed to remove 0.1 gaps Trimal (version v1.4.rev15) [56]. They were concatenated together using Geneious Prime 2025.0[57]. The tree was built using IQ-TREE (version 1.6.12), ultrafast bootstraps, and model finder plus. For ease of interpretation, we separated the tree into six miniature trees based on phylum phylogeny (**Supplemental Figure 2**). We selected the six based on abundance and lifestyle implications. References were chosen using BlastP and GTDB-TK [52].

### Ecosystem comparisons with genomes and rpS3

To understand how closely related genomes were in the ecosystem, we compared genomes using dRep, at increasing levels of 97%, 98%, 99%, and 99.5% ANI, to identify if and when a genome of similar ANI was shared across sites. To understand how similar organisms were between the hyporheic zone and river datasets to the hillslope and floodplain, we pulled rpS3 genes using Aksha. While we focus on genomes, there are organisms that are not binned, so we used rpS3 after Spencer et al. 2019 [44]. To determine the presence of organismal overlap, we used rpS3 genes as a marker for taxonomy. We looked for matches using MMseqs clustering at 90%, 95%, and 97%. Given the few instances of a high percent of matches (>99%), we did not pursue an inStrain analysis at the genome level.

### Identification of strains using Instrain

Identical strains were identified using inStrain and were based on comparing read mapping to the same subspecies [58]. Subspecies were considered based on preliminary information from RPS3 matches and dereplication results. A genome was considered an identical strain in two samples if 99% of the contigs of the genome from both samples shared greater than 99.999% population-level ANI. The 99.999% popANI cutoff was based on a previously established threshold [58]. popANI considers both major and minor alleles as opposed to consensus average nucleotide identity, which is calculated on the consensus sequence alone. The strains that we saw in the sample were shared by EAQ-C and Shumway. We note that DNA from Shumway well was extracted and sequenced on a run separate from the EAQ-C samples, which eliminates concerns of contamination.

### Giant Proteins

Given the identification of two proteins of >30,000 amino acids, the genes for which were not terminated by a stop codon, we validated the assembly and used manual curation to extend the contigs. Internal validation required stringent support of the sequence by mapped reads (coverage by reads with ≤ 1% SNP relative to the reference). Extension was performed by mapping paired reads to the ends of the de novo assembled contigs. Small extensions achieved in each round of extension enabled recruitment of small (few kbp) *de novo* assembled contigs that were added to the contig ends if there was near-perfect or perfect contig overlap. Extended contigs were re-validated after each step. Given the low coverage in any sample, but the presence of contigs from the same Omnitrophota in many samples, we combined reads from the four samples in which the genome was at highest coverage (all from the EAQ C site). Contig extension was terminated when preceding genes were apparent and when no clear path from the gene end could be deciphered. Read recruitment and curation were performed in Geneious Prime 2025.0[57]. Domains on the giant proteins were identified with Pfam V38.1 [59].

### Identification of genes potentially involved in inter-organism competition

HMMs from DefenseFinder [60] were used to identify the presence of Type VI secretion systems, RHS toxins, T6SS effectors, and lytic proteins within a set of genomes dereplicated at 95%.

## Acknowledgements

We are grateful to Curtis Beutler and Alex Newman for transporting us to the Pumphouse site. We would like to thank Luis Valentin-Alvarado and Andrew Turner for their support sampling. This material is based partially upon work supported as part of the Watershed Function Scientific Focus Area funded by the U.S. Department of Energy, Office of Science, Office of Biological and Environmental Research under Contract No. DE-AC02-05CH11231.

We thank Rocky Mountain Biological Laboratory for the use of their lab space. We thank Rohan Schadeva for his technical support. We acknowledge the sequencing services provided by Maryland Genomics at the Institute for Genome Sciences, University of Maryland School of Medicine. We also acknowledge some sequencing was performed at QB3 Genomics, UC Berkeley, Berkeley, CA, RRID:SCR_022170.

## Author Contributions

SM and JFB designed the study with guidance on hydrological conditions from MEN, KHW, ELB. Susan performed the metagenomic assemblies. SM, JW-R, and JFB polished the draft genomes and JFB performed manual curation, with assistance from SM. SM analyzed the rpS3 marker genes, strain overlap, community composition, and lifestyles of organisms, with input from L-XC, JW-R, and JH. SM identified the giant proteins and JW-R analyzed the structure of the giant proteins. SL supported data management. SM, JW-R, and JFB drafted the figures. SM and JFB wrote the manuscript, with input from all authors. All authors finalized and approved the manuscript.

## Data availability

The genomes described in this project can be found on https://ggkbase.berkeley.edu/east_river_hyporheic_zone_analysis_drep_95/organisms.

## Supplementary Materials

Supplemental Figures 1-3

Supplemental Tables 1-3

