## Supplemental Figures for "Establishment of microbial strains from the river, groundwater, and soil in the hyporheic zone is limited despite connectivity"

**This PDF includes supplemental figures 1-3.**

**Other supplementary materials for this manuscript include supplemental tables 1-3.**

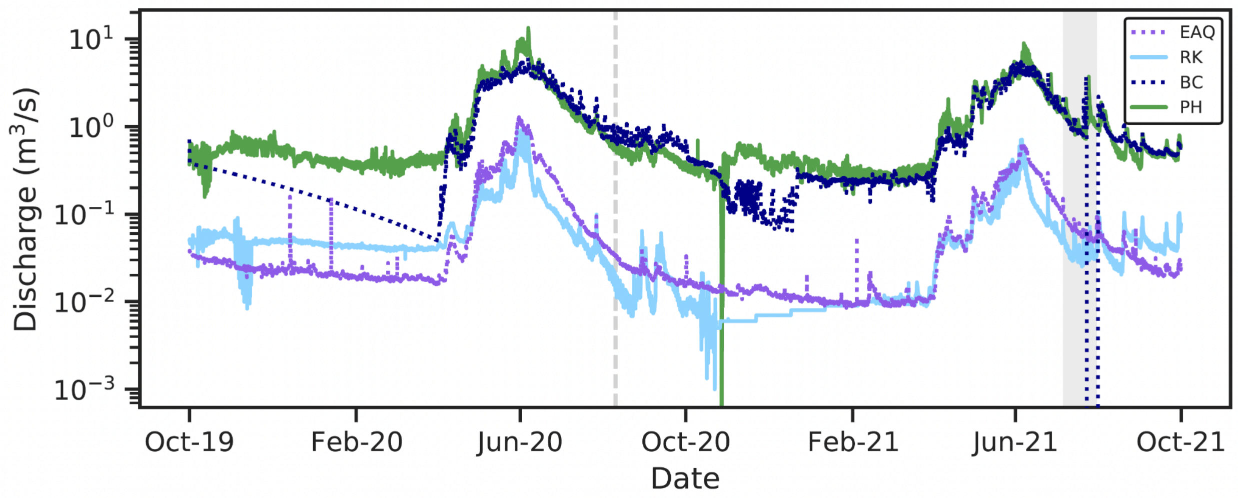


**Supplemental Fig. 1: Temporal discharges in log scale.** Discharge flow of the river water in the site over a two-year period in log scale. The dashed line indicates when samples were collected in the summer of 2020, and the shaded area shows the timeframe when we sampled in the summer of 2021.

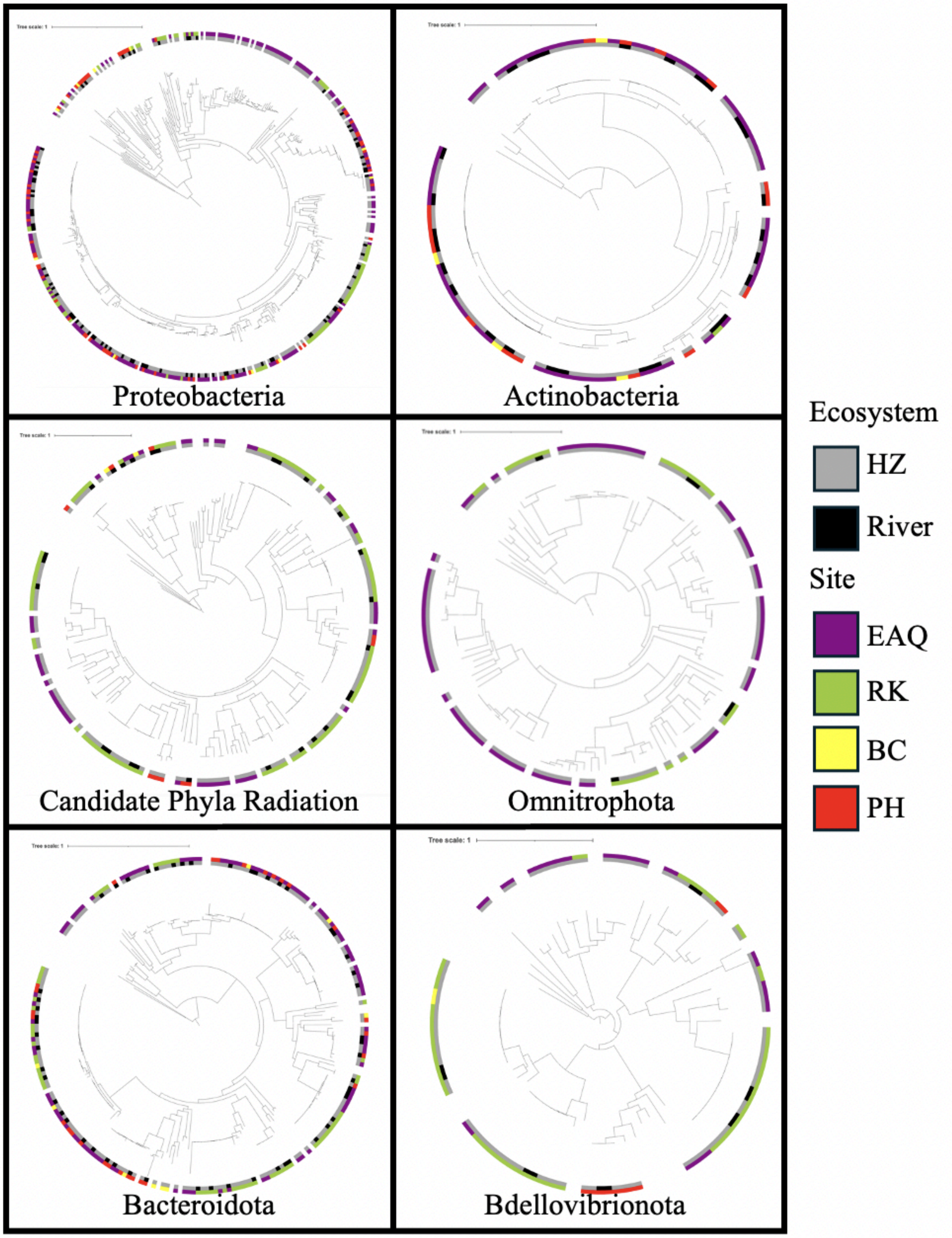


**Supplemental Figure 2: Diversity of six selected phyla along the river corridor in the hyporheic zone and river.** Trees of concatenated ribosomal proteins show to identify groupings of similar organisms. Color shows the location along the watershed and black and grey show the ecosystem compartment, river versus hyporheic zone. We selected the six phyla based on factors including lifestyle and relative abundance in the samples.

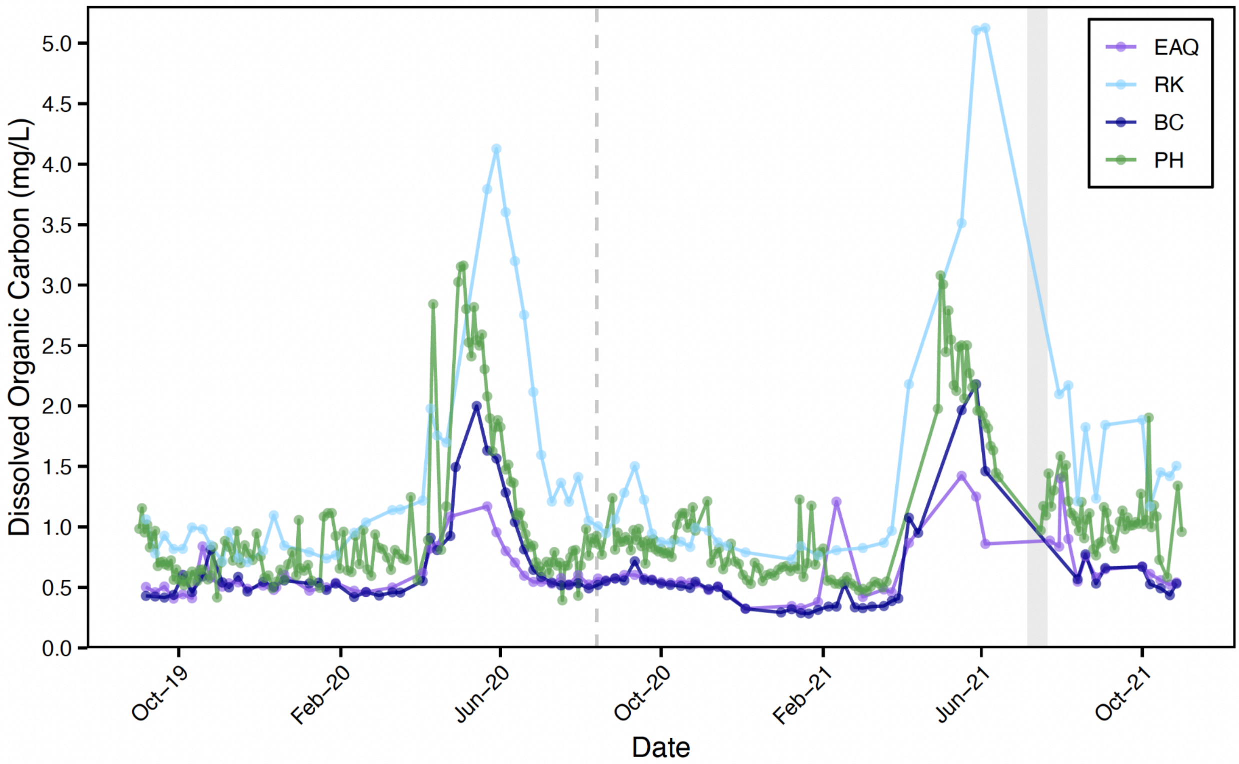


**Supplemental Figure 3**: **Dissolved organic carbon along the East River.** Dissolved organic carbon data is shown over a two-year period for EAQ, RK, BC, and PH. A grey dashed line indicates when samples were taken in 2020 and the shaded area shows the timeframe when they were sampled in 2021. The data for dissolved organic carbon  data is available at: doi:10.15485/1660459.
